# Neural Correlates of Working Memory Training: Evidence for Plasticity in Older Adults

**DOI:** 10.1101/869164

**Authors:** Alexandru D. Iordan, Katherine A. Cooke, Kyle D. Moored, Benjamin Katz, Martin Buschkuehl, Susanne M. Jaeggi, Thad A. Polk, Scott J. Peltier, John Jonides, Patricia A. Reuter-Lorenz

**Affiliations:** Department of Psychology, University of Michigan, 530 Church St, Ann Arbor, MI 48109, United States; Department of Mental Health, Bloomberg School of Public Health, Johns Hopkins University, 615 N Wolfe St, Baltimore, MD 21205, United States; Department of Human Development and Family Science, Virginia Tech, 295 W Campus Dr, Blacksburg, VA 24061, United States; MIND Research Institute, 111 Academy Dr, Irvine, CA 92617, United States; School of Education, University of California, Irvine, 3200 Education Bldg, Irvine, CA 92697, United States; Functional MRI Laboratory, Department of Biomedical Engineering, University of Michigan, 2360 Bonisteel Blvd, Ann Arbor, MI 48109, United States

**Author notes:** **Corresponding Authors:** Alexandru D. Iordan, Ph.D. & Patricia A. Reuter-Lorenz, Ph.D., 530 Church St, Ann Arbor, MI 48109, United States.

**Keywords:** executive functions, fronto-parietal, default-mode, cognitive training, aging

## Abstract

Brain activity typically increases with increasing working memory (WM) load, regardless of age, before reaching an apparent ceiling. However, older adults exhibit greater brain activity and reach ceiling at lower loads than younger adults, possibly reflecting compensation at lower loads and dysfunction at higher loads. We hypothesized that WM training would bolster neural efficiency, such that the activation peak would shift towards higher memory loads after training. Pre-training, older adults showed greater recruitment of the WM network than younger adults across all loads, with decline at the highest load. Ten days of adaptive training on a verbal WM task improved performance and led to greater brain responsiveness at higher loads for both groups. For older adults the activation peak shifted rightward towards higher loads. Finally, training increased task-related functional connectivity in older adults, both within the WM network and between this task-positive network and the task-negative/default-mode network. These results provide new evidence for functional plasticity with training in older adults and identify a potential signature of improvement at the neural level.

## Introduction

Working memory (WM) is a fundamental cognitive ability that typically declines with age (Park et al., 2002; Salthouse, 1994). Functional neuroimaging evidence indicates age differences in neural recruitment (Li et al., 2015; Spreng, Wojtowicz, & Grady, 2010) and suggests that WM load influences whether older adults will over-activate or under-activate WM circuitry relative to younger adults (Cappell, Gmeindl, & Reuter-Lorenz, 2010; Heinzel et al., 2014; Schneider-Garces et al., 2010). In particular, older adults tend to over-activate PFC regions at lower loads, while performing equivalently to younger adults, but under-activate at higher loads, while performing more poorly than younger adults (Cappell et al., 2010).

According to the Compensation Related Utilization of Neural Circuits Hypothesis (CRUNCH) (Reuter-Lorenz & Cappell, 2008) (see also Cabeza et al., 2018), over-activation in older adults compensates for age-related decline in neural efficiency. While lower levels of task demand can be met by over-recruitment, increased demand will rapidly outstrip resource availability in older adults, resulting in a performance drop and activation decrease (Cappell et al., 2010; Mattay et al., 2006). The inflection point along the task-demand axis where activity reaches its peak level and then declines (i.e., the crunch point) is thought to reflect a resource ceiling beyond which neural mechanisms are inefficiently engaged (Fig. 1a). Older adults reach this apparent resource ceiling at lower loads than younger adults (Cappell et al., 2010), although younger adults can also show compensatory neural activation when task demands are sufficiently high (Holler-Wallscheid, Thier, Pomper, & Lindner, 2017).

**Fig. 1.**
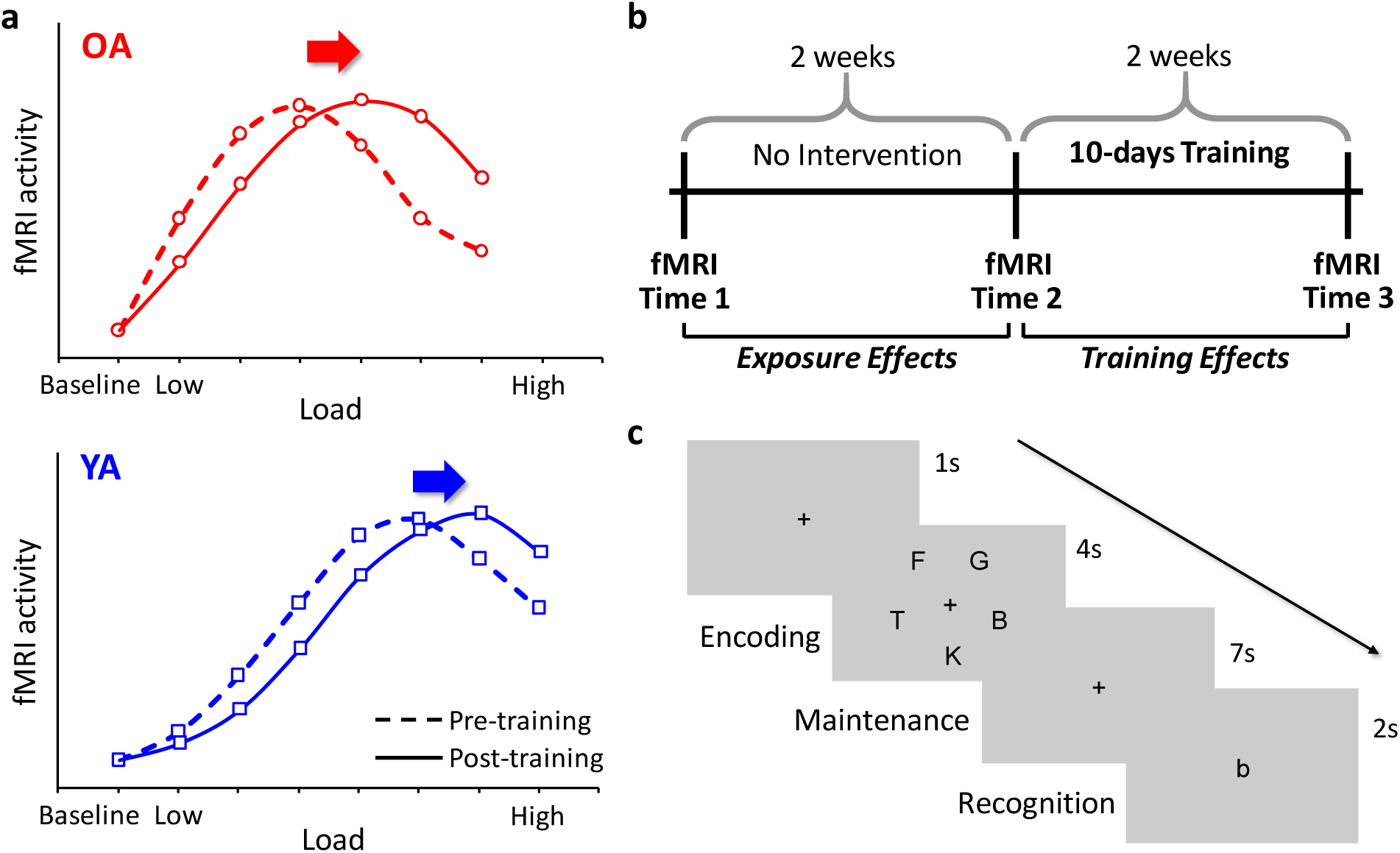
CRUNCH predictions and current design. **a**, CRUNCH predicts a rightward shift of the neural recruitment curves with training, regardless of age. **b**, The present within-subjects design enabled the dissociation of *task-exposure* (Time1 vs. Time2) from *training* (Time2 vs. Time3) effects. **c**, During each fMRI session, participants performed a delayed match-to-sample verbal WM task, with varying memory sets. OA, older adults; YA, younger adults.

While accumulating evidence clearly supports the idea of load-dependent over-activation in older adults, particularly when able to reach performance levels comparable with younger adults (e.g., Berlingeri, Danelli, Bottini, Sberna, & Paulesu, 2013; Cappell et al., 2010; Heinzel et al., 2014; Holler-Wallscheid et al., 2017; Kennedy et al., 2015; Schneider-Garces et al., 2010) (for a recent meta-analysis see Li et al., 2015), the hypothesis that over-activation signals compensation has sometimes been challenged. It could be argued that the correlational nature of the available evidence makes it impossible to ascertain whether increased brain activity is used in the service of improved performance (Schneider-Garces et al., 2010). Alternatively, a common factor, such as general insufficient capacity or deterioration of neural function (e.g., due to insult or aging), could be at the root of both increased activity and decreased capacity or performance (Morcom & Henson, 2018; Spreng et al., 2010).

Posing this problem in an intervention framework (i.e., training study) provides longitudinal data to complement existing correlational findings and using a within-subjects design accounts for potential confounding by a common general factor. CRUNCH makes clear predictions about how activation in regions critical to WM should change due to training (Lustig, Shah, Seidler, & Reuter-Lorenz, 2009). Specifically, training should simultaneously (1) reduce activation under low demand, consistent with the idea of reduced need for compensatory over-activation with training, and (2) increase activation under high demand, consistent with the idea of enhanced dynamic range of activation (i.e., greater responsivity under high demand) with training (Kennedy, Boylan, Rieck, Foster, & Rodrigue, 2017). In other words, CRUNCH predicts a rightward shift of the demand-activation curve with training, irrespective of age (Fig. 1a).

The main goal of the present study was to test this hypothesis in the context of a within-subjects intervention design. Achieving this goal would further shed light on potential neural mechanisms of plasticity mobilized by cognitive training. Such mechanisms have heretofore been characterized mainly by decreases in activation, particularly in the WM network (Bamidis et al., 2014; Belleville & Bherer, 2012; Bherer, 2015; Brehmer, Kalpouzos, Wenger, & Lovden, 2014; Lustig et al., 2009). To the extent that these mechanisms relate to CRUNCH, training should also lead to increased activation in WM regions at high WM loads.

To summarize, if a successful training intervention simultaneously (1) improves WM performance and (2) shifts the demand-activation curve towards higher loads within the same neural circuits that showed over-activation before training, then it would provide strong evidence for the compensatory nature of neural over-activation. We tested this hypothesis in a sample comprising both healthy older and younger adults, who participated in an adaptive verbal WM training study with 3 functional MRI scanning sessions (see Fig. 1b). Sessions 1 and 2 were two weeks apart (Time1 and Time2) and preceded a 10-day adaptive WM training intervention. The third scanning session (Time3) was conducted immediately after training, approximately two weeks after Time2.

Based on *a priori* considerations (see Cabeza et al., 2018; Lustig et al., 2009), our approach comprised 3 analytic components: First, to dissociate the effects of *task-exposure* from the effects of *training*, we performed two comparisons, specifically Time1 vs. Time2 for task-exposure effects and Time2 vs. Time3 for training effects. Second, to account for the possibility that older adults may recruit additional brain regions compared to younger adults, we assessed both task-exposure and training effects *between* groups, using meta-analytically defined regions, and training effects *within* each group, using age-specific maps independently identified at Time1. Finally, to determine whether the CRUNCH model and its implications for mechanisms of training could be further extended to measures of functional coupling between regions critical to WM performance, analyses of brain activation were supplemented by analyses of functional connectivity.

## Materials and Methods

### Participants

A sample of 23 healthy, cognitively normal older and 23 younger adults was recruited from the University of Michigan campus and community surrounding Ann Arbor, Michigan. Initial sample size was based on prior work examining age and load effects in WM (Cappell et al., 2010). All participants were right-handed, native English speakers with normal or corrected-to-normal hearing and vision and were screened for history of head injury, psychiatric illness, or alcohol/drug abuse. Data from 2 older and 2 younger adults were excluded due to technical errors in the administration of the training (1 older adult) or fMRI (1 younger adult) tasks, inability to perform the fMRI task (1 younger adult did not provide responses to >50% of the trials), and attrition (1 older adult failed to return for the last scan). Thus, the sample for behavioral analyses consisted of 21 older adults (age range: 63-75; 10 women) with a mean age of 67.81 (± 3.31) years and 21 younger adults (age range: 18-28; 12 women) with a mean age (±S.D.) of 21.33 (± 2.65) years (see Table 1). In addition, 2 older adults were excluded from the fMRI analyses due to technical issues related to brain-imaging data acquisition, and thus the fMRI sample consisted of 19 older and 21 younger adults. Older adult participants completed the Short Blessed Test (Katzman et al., 1983) over the phone prior to inclusion in the study to screen for potential mild cognitive impairment, and additional neuropsychological assessments using the Montreal Cognitive Assessment (Nasreddine et al., 2005) confirmed normal cognitive function for all participants (scores ≥ 25). Additionally, participants were screened for depressive symptoms that could affect cognitive functioning using the depression module of the Patient Health Questionnaire (Kroenke, Spitzer, & Williams, 2001). Additional neuropsychological testing was performed at each assessment time point (Times1-3) and this is reported elsewhere (see Moored et al., in prep.). The University of Michigan Institutional Review Board approved all procedures, and all participants provided informed consent prior to participating.

**Table 1.**
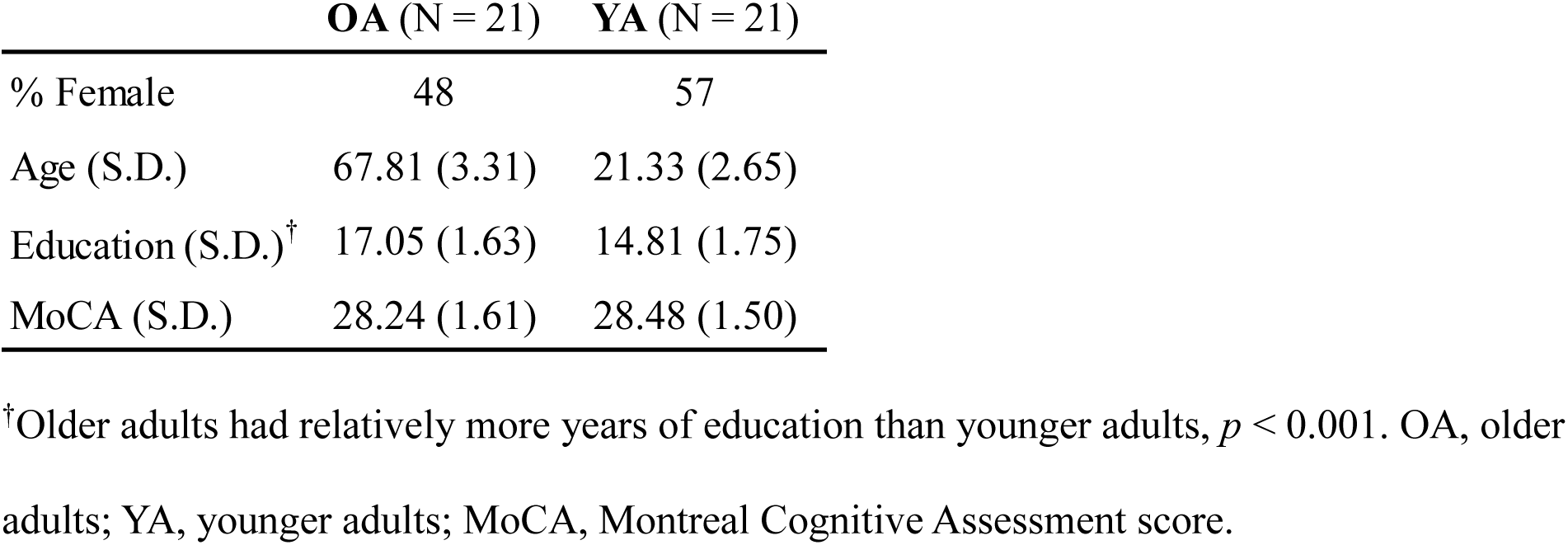
Sample demographic information.

### Experimental Design and Procedure

#### fMRI WM Task

During each of the 3 fMRI scanning sessions, participants performed a delayed match-to-sample verbal WM task (Sternberg, 1966) with span and supraspan loads (Fig. 1c). At the beginning of each trial, a set of letters was displayed during encoding (4 s), followed by a fixation cross during the maintenance interval (7 s). At retrieval, a probe letter was displayed on the screen (2 s), and participants indicated by a button-press whether or not the probe was part of the memory set. The memory sets varied in size from 4 to 8 letters for older adults and from 5 to 9 letters for younger adults. These age-specific ranges of loads were chosen based on pilot data to minimize ceiling and floor effects on WM performance, and to allow comparisons of both baseline performance and training-induced improvement. Both groups also completed a control condition (set size of 1) that served as an active baseline for the fMRI analyses (see below). During each fMRI session, participants completed 6 blocks of 24 trials, with each block comprising 4 trials of each set size, displayed in random order.

#### Behavioral WM Training Task

The training task was an adaptive verbal WM task, similar to the fMRI task in terms of the type of stimuli employed (i.e., letters) but different with respect to the set sizes and timing, as described below (Iordan et al., 2018). All participants started the first training session with a set size of 3 letters. The number of letters in each memory set remained constant for each block and was determined by the participant’s performance in the previous block. The set size increased by one letter if the participants’ accuracy was >86% on the preceding block and decreased by one letter if their accuracy was <72%. The set size attained in the last session of each day was used as the starting set size the subsequent day. For each trial, the memory set was displayed for a duration weighted by its size (325 ms × set size) at encoding, followed by a 3 s maintenance interval, and a 2 s retrieval period. Participants completed 6 blocks of 14 trials during each of the 10 training sessions. Both tasks were presented using E-Prime 2.0 (Psychology Software Tools, Pittsburgh, PA).

### Imaging Protocol

Imaging data were collected using a 3 T General Electric MR750 scanner with an eight-channel head coil. Functional images were acquired in ascending order using a spiral-in sequence, with MR parameters: TR = 2000 ms; TE = 30 ms; flip angle = 90°; field of view = 220×220 mm^2^; matrix size = 64×64; slice thickness = 3 mm, no gap; 43 slices; voxel size = 3.44×3.44×3 mm^3^. After an initial ten seconds of signal stabilization, 168 volumes were acquired for each of the 6 runs. A high-resolution T_1_-weighted anatomical image was also collected following the WM task, using spoiled-gradient-recalled acquisition (SPGR) in steady-state imaging (TR = 12.24 ms, TE = 5.18 ms; flip angle = 15°, field of view = 256×256 mm^2^, matrix size = 256×256; slice thickness = 1 mm; 156 slices; voxel size = 1×1×1 mm^3^). Images were produced using a k-space de-spiking of outliers, followed by reconstructing using an in-house iterative reconstruction algorithm with field-map correction (Sutton, Noll, & Fessler, 2003), which has superior reconstruction quality compared to non-iterative conjugate phase reconstruction. Initial images and field-map estimates were inspected for distortions and when present, the field maps were re-estimated using maps from adjacent runs.

### Behavioral Data Analyses

Responses in the WM task were classified in one of the four categories derived from signal detection theory (Green & Swets, 1966; Macmillan & Creelman, 2005): (1) *Hits*, corresponding to letters in the memory set correctly classified as Old, (2) *Misses*, corresponding to letters in the memory set incorrectly classified as New, (3) *Correct Rejections* (CRs), corresponding to new letters correctly classified as New, and (4) *False Alarms* (FAs), corresponding to new letters incorrectly classified as Old. Average percentages of probes correctly identified as being *Old* or *New* were also calculated for each participant [%WM Accuracy = (%Hits + %CR)/2], separately for each load and time point. %WM Accuracy is mathematically equivalent to %Corrected Recognition (%Hits – %False Alarms) and is typically used in WM studies with a neuroimaging component to capitalize on all possible trials (see below).

To investigate *task-exposure* effects, we compared accuracy for the two time points before training (i.e., Time1 vs. Time2), whereas to investigate *training* effects, we compared accuracy pre-vs. post-training (i.e., Time2 vs. Time3). We used mixed-design ANOVAs with Group (older vs. younger adults) as a between-subjects factor and Time (Time1 vs. Time2/ Time2 vs. Time3) and Load (loads 5-8) as within-subject factors. Of note, only loads 5-8 were used for between-subjects comparisons because these were included in the tasks for both older and younger adults, whereas set size 4 was unique to older adults and set size 9 was unique to younger adults.

Statistical analyses were performed using SPSS 24 (IBM Corp., Armonk, NY). A Greenhouse-Geisser correction for violation of sphericity was applied as needed, for all ANOVA models. Effect-sizes are reported as partial eta squared (η_p_^2^). At Time1, WM performance for 2 participants (1 older and 1 younger adult) was calculated based on 5 runs, due to technical issues occurring during 1 run, and response buttons were remapped for an older adult who switched the buttons for the first 3 runs; fMRI analyses matched these adjusted behavioral data.

### fMRI Data Analyses

Statistical analyses were performed using SPM12 (Wellcome Department of Cognitive Neurology, London) and MATLAB R2015a (The MathWorks Inc., Natick, MA), and were preceded by several preprocessing steps. Functional images were slice-time corrected, realigned, and co-registered to the anatomical image using a mean functional image. A study-specific anatomical template was created (younger and older adults together) (Iordan et al., 2018), using Diffeomorphic Anatomical Registration Through Exponentiated Lie Algebra (DARTEL) (Ashburner, 2007), based on segmented grey matter and white matter tissue classes, to optimize inter-participant alignment (Klein et al., 2009). The DARTEL flowfields and MNI transformation were then applied to the functional images, and the functional images were resampled to 3×3×3 mm^3^ voxel size and smoothed (8 mm full width at half maximum Gaussian kernel). The average proportion of outlier volumes (differential motion *d* > 3 mm or global intensity *z* > 6, identified with Artifact Detection Toolbox [ART]; www.nitrc.org/projects/artifact_detect) was <.5% (older adults: 0.28%; younger adults: 0.48%), and there were no significant differences in the number of outlier volumes or maximum motion between the two groups at any time point (Times1-3) or within-groups across time points (Time1 vs. Time2 or Time2 vs. Time3), as assessed by permutation testing (10^5^ permutations, two-sided, all *p*s > .05).

At the first level, each participant’s preprocessed functional data were analyzed using an event-related design in the general linear model (GLM). The GLMs included separate regressors for each load (1, 4-8 for older adults/5-9 for younger adults) at each task phase (encoding, delay, and probe), resulting in 18 regressors. In addition, the model included 1 regressor for incorrectly answered trials and 6 regressors for the realignment parameters derived from preprocessing (3 translations and 3 rotations). To further reduce residual influence of motion that may not be properly explained by the realignment parameters, scan nulling regressors were added for any outlier volumes (i.e., 1 for the outlier volume and 0 everywhere else) identified using the procedure described above; this avoids discarding entire datasets and is mathematically equivalent to extracting outlier volumes while preserving temporal continuity (Caballero-Gaudes & Reynolds, 2017; Lemieux, Salek-Haddadi, Lund, Laufs, & Carmichael, 2007; Whitfield-Gabrieli et al., 2011).

Evoked hemodynamic responses to all events were modeled with a delta (stick) function corresponding to the onset of each event convolved with a canonical hemodynamic response function, in conjunction with a high-pass filter (128 s) and an intensity threshold of 70% (to avoid inclusion of regions susceptible to fMRI signal drop-out) (Iordan et al., 2018), and runs were modeled separately. To differentiate maintenance-related activity for the different loads, analyses were restricted to the task-delay phase, and linear contrasts were defined for each load (older adults: loads 4-8; younger adults: loads 5-9) relative to the load of 1 (active baseline).

### Brain Activation Analyses

#### Analyses of Brain Activity in Meta-analytically Defined WM Regions

Similar to the behavioral analyses, we investigated both task-exposure (Time1 vs. Time2) and training effects (Time2 vs. Time3) on brain activity. To identify brain regions associated with WM processing, we used an independently-defined functional mask derived by meta-analysis performed with Neurosynth (Yarkoni, Poldrack, Nichols, Van Essen, & Wager, 2011). The map was derived by an automated meta-analysis performed on studies indexed by the feature “working memory” (1091 studies [accessed October 5^th^, 2018], reverse inference/association test map, thresholded at *p*_FDR_ < 0.01, which is the lowest default threshold in Neurosynth; see Fig. S1a). The binarized Neurosynth WM mask (resampled to 3×3×3 mm^3^ voxel size) was used for region of interest (ROI) analyses using MarsBaR (Brett, Anton, Valabregue, & Poline, 2002). The GLM described above was run on the time-course of average activity within the ROI and the resulting contrast values for each participant, time point, and load were exported to SPSS and analyzed with mixed-design ANOVAs (Group×Time×Load).

Finally, to test for a rightward shift in the load-dependent neural recruitment (CRUNCH) curve with training, we ran multilevel models on brain activity with loads nested within participants and a random intercept for Load, separately for each group, before (Time2) and after training (Time3). A rightward shift of the CRUNCH curve would be consistent with a model comprising an additional quadratic term (Load^2^) describing the data over and above a model comprising only a linear term (Load) before but not after training. We employed full maximum-likelihood estimation, which allows model comparison, and the change in model fit (i.e., difference between log-likelihoods, *-2LL*) from linear to quadratic was assessed using a chi-square test (χ^2^) with 1 degree of freedom.

#### Analyses of Brain Activity in Group-Specific Regions Sensitive to Load Manipulation

To identify group-specific regions sensitive to the load manipulation, brain-wide contrast images generated for each participant at Time1 were analyzed at the second level, separately for each group. First, a voxel-wise one-way repeated-measures ANOVA with Load as factor (*F*-contrast: [1 −1 0 0 0; 0 1 −1 0 0; 0 0 1 −1 0; 0 0 0 1 −1]) identified brain regions showing up- *or* down-regulation in response to the Load manipulation (i.e., up-regulation is typically expected in fronto-parietal regions, whereas down-regulation is typically expected in default-mode regions, for high relative to low WM loads (Ceko et al., 2015; Shulman et al., 1997); see Fig. S1b). Then, task-positive and task-negative brain regions were separated by masking (logical “AND” conjunction) (Nichols, Brett, Andersson, Wager, & Poline, 2005) the *F*-maps with binarized composite random-effects *t*-maps identifying up- and down-regulation, respectively, in response to the load manipulation. Specifically, the composite *t*-map identifying regions showing *up*-regulation was constructed by masking (logical “OR” conjunction) the binarized group-level *t*-maps derived by testing for effects of each load > baseline (one-sided). Similarly, the composite *t*-map identifying regions showing *down*-regulation was constructed by masking (logical “OR” conjunction) the binarized group-level *t*-maps derived by testing for effects of baseline > each load (one-sided). Unless specified otherwise, we used a cluster-forming threshold of *p* < 0.001, in conjunction with a Random Field Theory familywise error (FWE) correction of *p*_FWE_ < 0.05 (Eklund, Nichols, & Knutsson, 2016; Flandin & Friston, 2017). These final, group-specific masks independently defined at Time1 were then employed as ROIs to analyze brain activity pre-vs. post-training (Time2 vs. Time3), using MarsBaR.

### Functional Connectivity Analyses

#### Analyses of Functional Connectivity within the Meta-analytically Defined WM Network

To complement the brain activation analyses described above, we also investigated task-exposure and training effects on functional connectivity within the WM network. To identify the WM network, we employed the (Power et al., 2011) functional atlas in conjunction with the meta-analytical WM mask described above. We chose the Power et al. atlas because it comprises cortical, subcortical, and cerebellar ROIs derived meta-analytically across a variety of tasks. First, 5 mm-radius spheres were centered at each of the 264 coordinates of the atlas. Then we selected those ROIs that had at least 8 voxels (∼50% volume) overlap with the meta-analytical WM map (see above), thus retaining 23 ROIs (see Fig. S3a).

Functional connectivity between the selected ROIs was calculated using correlational psychophysiological interaction (cPPI) analysis, which computes the partial correlation between PPI terms of any two ROIs while controlling for effects of co-activation, task-unrelated coupling, and nuisance signals (Fornito, Harrison, Zalesky, & Simons, 2012). To parallel the univariate analyses, we focused on the delay activity and used task regressors based on the GLM described above. Time courses for each ROI (extracted using REX, Whitfield-Gabrieli & Nieto-Castanon, 2012) were first deconvolved, then multiplied by the unconvolved task regressor modeling the effect of each load vs. baseline (load of 1), and then reconvolved with the hemodynamic response function to generate the ROI-specific PPI terms. Task-related functional connectivity between any two ROIs was estimated as the partial correlation between those regions’ PPI terms, separately for each load, adjusted for the original task regressor (to control for co-activation effects), the original regions’ time courses (to control for task-unrelated coupling), the regressor for incorrect trials (from the GLM model), and nuisance signals (6 realignment parameters, regressors for outlier volumes, and average time-courses within white matter and cerebrospinal fluid masks). Correlation coefficients were then Fisher-z transformed to allow statistical testing. Within-WM network functional connectivity was estimated as the sum of all connectivity values divided by the number of possible connections (Geerligs, Renken, Saliasi, Maurits, & Lorist, 2015). Task-exposure and training effects were analyzed using Group×Time×Load ANOVAs, similar to previous analyses.

#### Analyses of Functional Connectivity in Group-Specific Networks Sensitive to Load Manipulation

Similar to the univariate analyses, we identified group-specific networks sensitive to the load manipulation based on Time1 results. Specifically, the task-positive and task-negative networks were identified based on the peak-activation voxels of the task-positive and task-negative *F*-maps resulting from the voxel-wise ANOVAs described above (see Fig. S3b and Tables 2 and 3 for employed coordinates). Then, 5 mm-radius spheres were centered at each Time1 peak coordinate, and functional connectivity between ROIs was calculated for Time2 and Time3, using cPPI, and the resulting correlation coefficients were Fisher-z transformed. Functional connectivity within the task-positive and task-negative networks, as well as between the two networks, was calculated in the same way as described above.

**Table 2.**
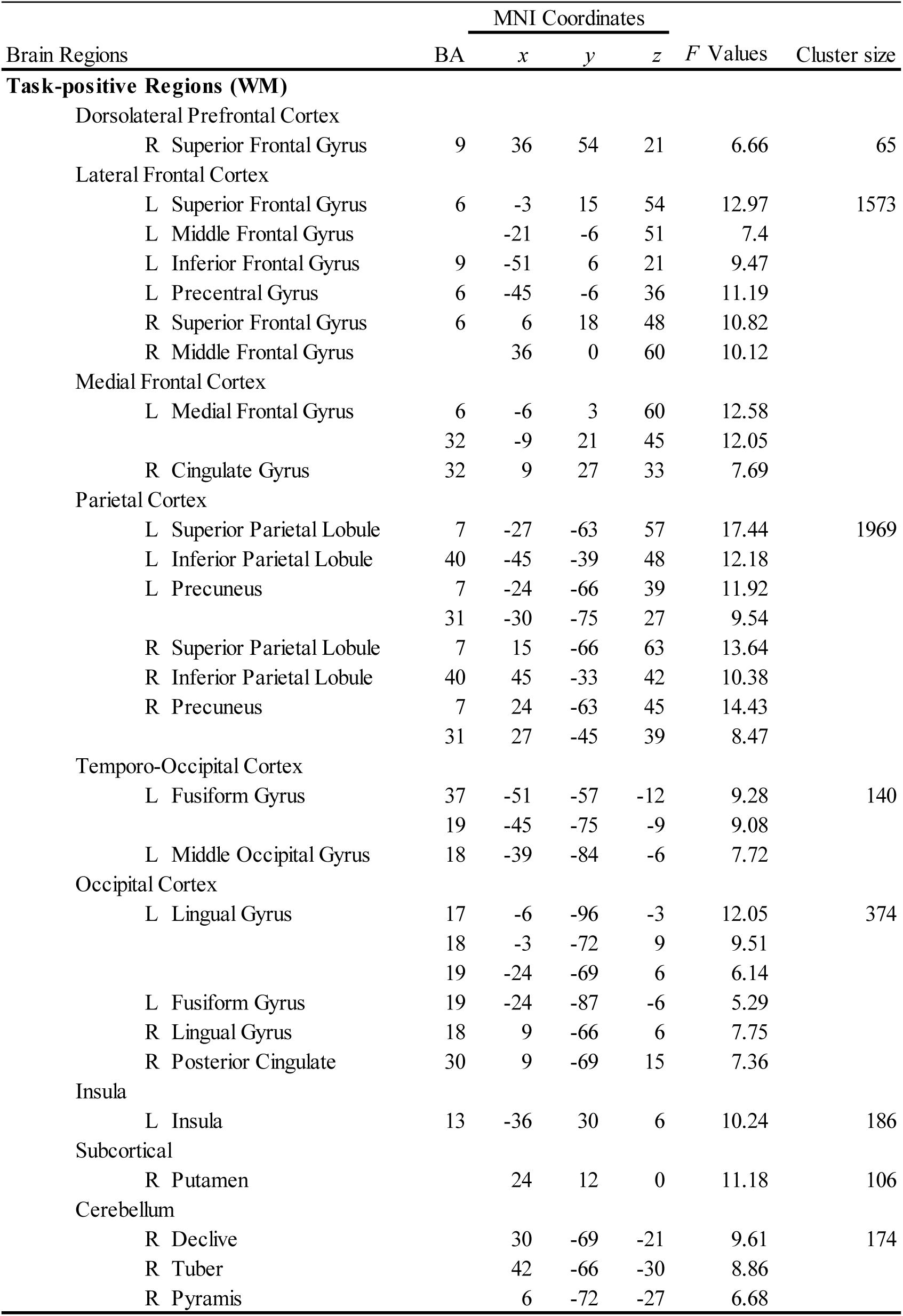
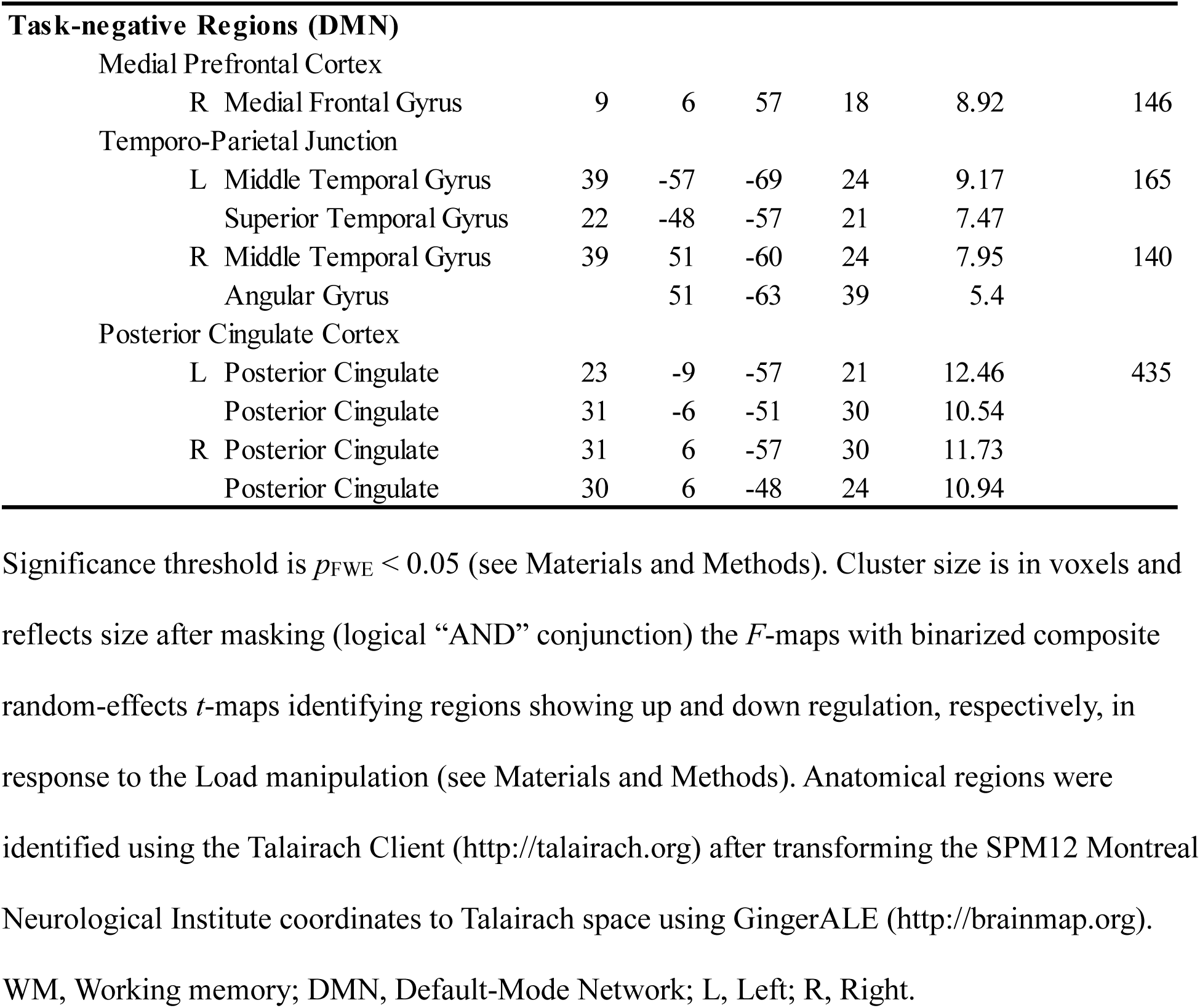
Task-positive and task-negative brain regions showing up and down regulation, respectively, in response to the Load manipulation, in older adults.

**Table 3.**
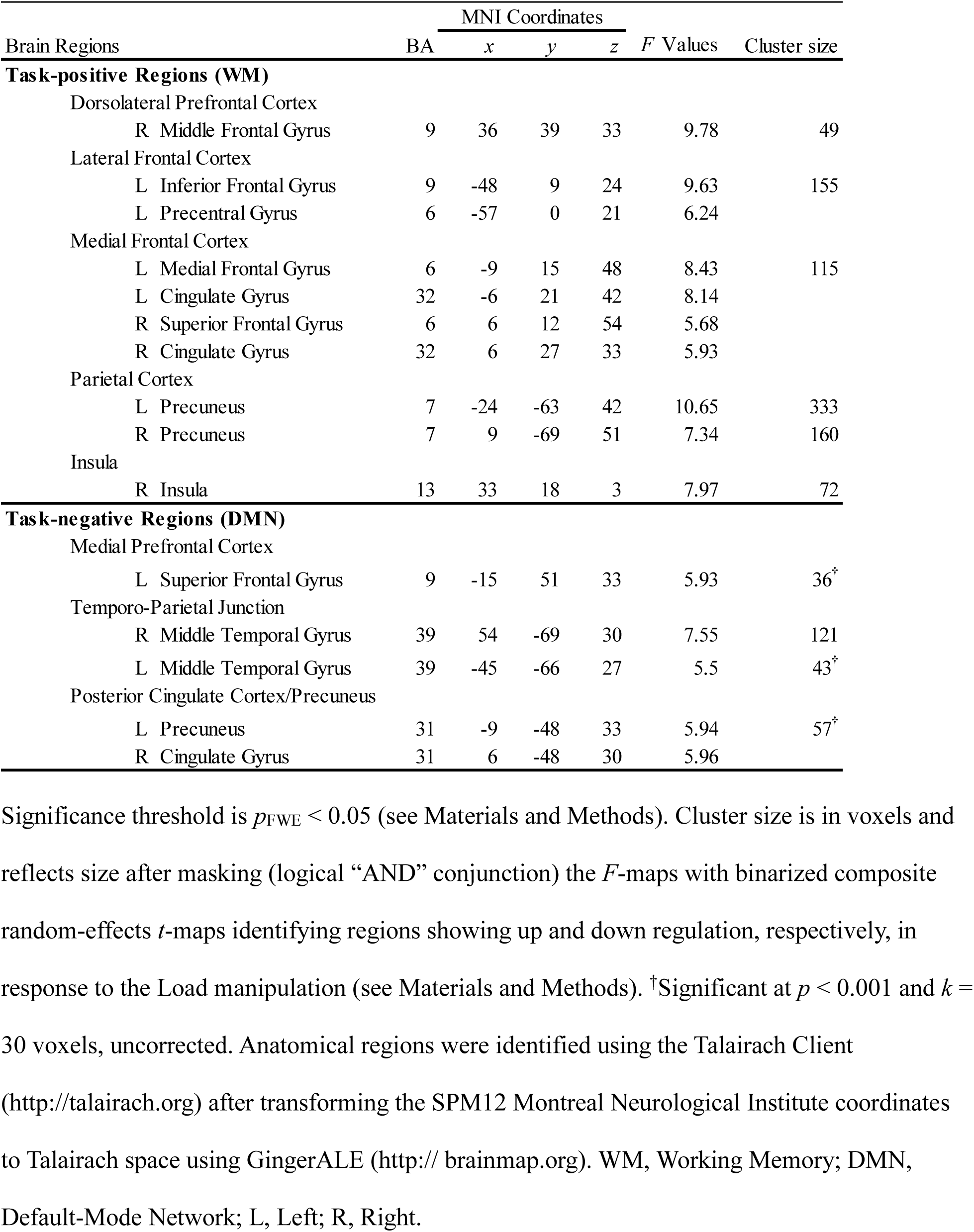
Task-positive and task-negative brain regions showing up and down regulation, respectively, in response to the Load manipulation, in younger adults.

## Results

### Behavioral Results

Effects of task-exposure and training on WM performance were examined with loads 5-8, which were common to both groups, using Group×Time×Load ANOVAs. The main effect of Load was significant at *p*<0.001 for all ANOVA models, unless noted otherwise. First, the task-exposure (Time1 vs. Time2) analysis (Fig. 2, left panel) showed that, while younger adults performed overall better than older adults (*F*_1,40_=5.91, *p*=0.02, η_p_^2^=0.13), this group difference was reduced with task exposure (Time×Group: *F*_1,40_=6.17, *p*=0.017, η_p_^2^=0.13). Second, analysis of training effects (Time2 vs. Time3) (Fig. 2, right panel) showed that performance improved with training for both groups (Time: *F*_1,40_=13.04, *p*=0.001, η_p_^2^=0.25).

**Fig 2.**
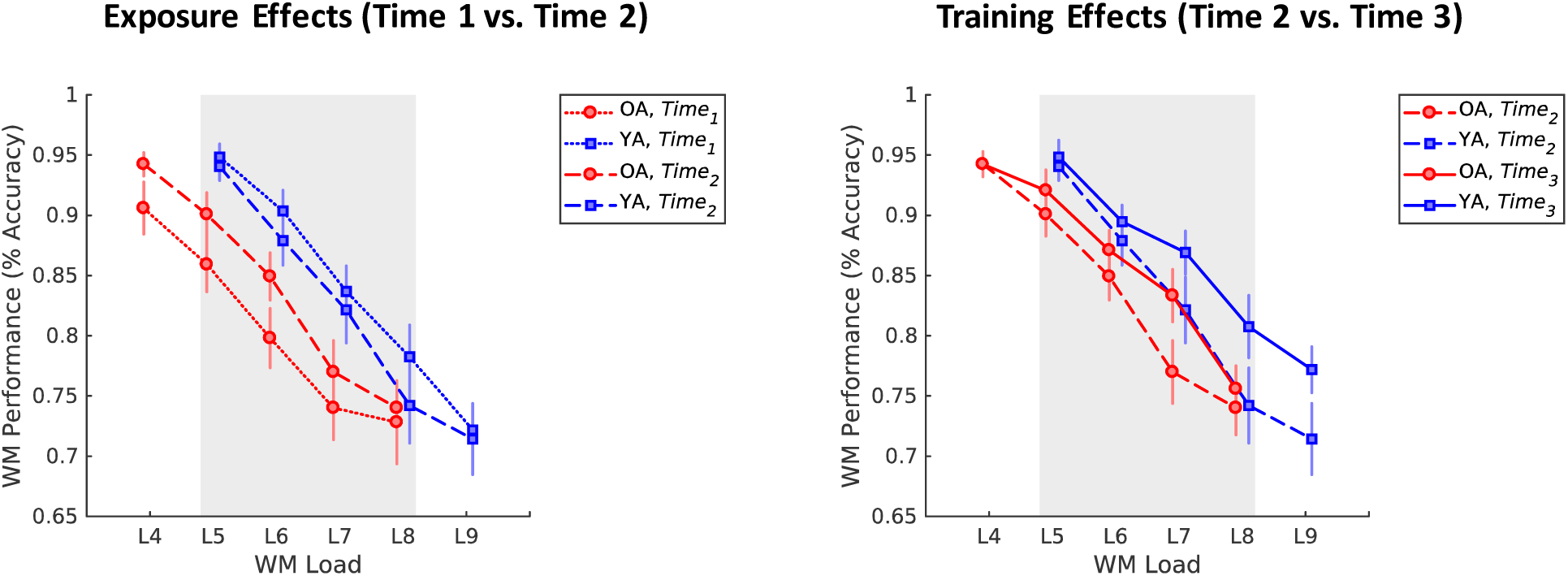
Exposure and training effects for WM performance. Behavioral results show a reduction in age-related differences in WM performance due to task exposure (left panel) and training-related improvements in both groups (right panel). Line-graphs display average WM performance for each load. Error bars display standard error of the mean. The grey rectangle highlights the loads common to both groups (loads 5-8). WM, working memory; OA, older adults; YA, younger adults.

### fMRI Results

We examined both activation in and functional connectivity between brain regions involved in WM performance. In each case, we performed two sets of complementary analyses, examining (1) task-exposure and training effects *between* groups, using meta-analytical WM maps/regions, and (2) training effects *within* each group, using age-specific maps/regions, independently identified at Time1. These latter analyses take into account the possibility that older adults may recruit additional brain regions compared to younger adults. The two complementary analyses enabled us to compare the age groups directly, using the same independently defined map/regions, and to assess how training affects the amplitude of brain activation and functional connectivity within groups, independent of age differences in the extent of activation. Similar to the behavioral analyses, the main effect of Load was significant at *p*<0.001 for all ANOVA models, unless noted otherwise

### Brain Activation Results

#### Task-exposure and Training Effects in Meta-analytically Defined WM Regions

First, we examined task-exposure and training effects on brain activity in WM regions identified meta-analytically using Neurosynth (Yarkoni et al., 2011) (see Materials and Methods and Fig. S1a). Paralleling the behavioral analyses, we performed two Group×Time×Load ANOVAs on estimates of average activity (loads 5-8) within this map of WM regions. The Time1 vs. Time2 analysis (Fig. 3, left panel) indicated no effect of task exposure (*F*_1,38_=0.81, *p*=0.373, η_p_^2^=0.021) and greater overall recruitment in older adults (*F*_1,38_=5.43, *p*=0.025, η_p_^2^=0.13), with a decline in activation at the highest load. Comparing Time2 and Time3 to assess training effects (Fig. 3, right panel) again indicated greater overall recruitment in older adults (*F*_1,38_=4.22, *p*=0.047, η_p_^2^=0.10), but also increased recruitment at higher loads with training for both groups (Time×Load: *F*_3,114_=3.54, *p*=0.017, η_p_^2^=0.09). Follow-up analysis confirmed that, with training, activation increased, rather than decreased, from load 7 to 8 in older adults (Time×Load interaction: *F*_1,18_=7.29, *p*=0.015, η_p_^2^=0.29) (see Supplementary Results for additional robustness analysis).

**Fig 3.**
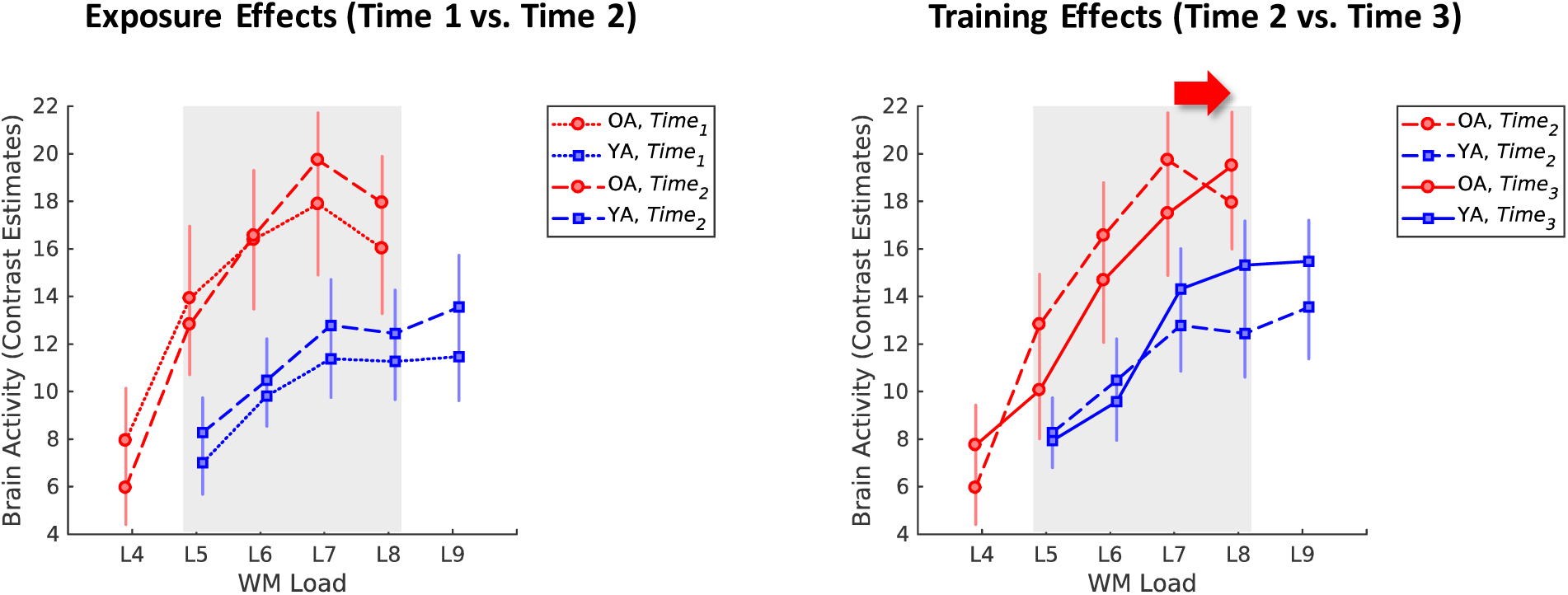
Exposure and training effects for brain activity in meta-analytically-defined WM regions. Brain imaging results show greater overall recruitment of WM regions in older adults pre-training (left panel) and increased activation at higher loads for both groups post-training (right panel), consistent with increasing responsiveness in the WM network. For older adults, this entails greater activation at the highest load, which is characterized by a shift from quadratic to linear trends (red arrow; see Results). For younger adults, greater responsiveness is evident across a range of higher loads. Line-graphs display average brain activity for each load vs. baseline (load of 1) in meta-analytically-defined WM regions (see Materials and Methods and Fig. S1a). Error bars display standard error of the mean. The grey rectangle highlights the loads common to both groups (loads 5-8). WM, working memory; OA, older adults; YA, younger adults.

Multilevel models on brain activity by Load (see Materials and Methods) confirmed a rightward shift of the CRUNCH curve with training that was specific to older adults: A quadratic trend described the data over and above a linear trend before (*-2LL*_linear_=473.81, *- 2LL*_linear+quadratic_=462.57, *χ*^2^_difference(1)_ =11.24, *p*<0.001) but not after training (*-2LL*_linear_=479.09, *- 2LL*_linear+quadratic_=476.74, *χ*^2^_difference(1)_ =2.35, *p*=0.126). For younger adults, quadratic terms did not significantly improve model fit (*p*s>0.17).

#### Training Effects in Group-specific Regions Sensitive to Load Manipulation

In the second set of analyses, group-specific maps sensitive to WM load were defined based on Time1 data and then training-induced changes (Time2 vs. Time3) were examined in each age group (see Materials and Methods). First, voxel-wise ANOVAs of brain activity at Time1, across all loads (i.e., loads 4-8 in older adults and 5-9 in younger adults), identified load-sensitive regions, separately for each group. These included both task-positive and task-negative regions, which overlap with canonical WM and default-mode network (DMN) regions, respectively (Fig. S1b and Tables 2 and 3). Then, training effects were analyzed separately for each group, using Time×Load ANOVAs on estimates of average activity within the task-positive and task-negative maps (see Materials and Methods).

In task-positive regions, both groups showed greater activation at higher loads after training compared to before training (Time×Load older adults: *F*_4,72_=3.48, *p*=0.012, η_p_^2^=0.16; younger adults: *F*_4,80_=3.14, *p*=0.019, η_p_^2^=0.14), thus replicating the Neurosynth results (see Fig. 4a; see also Supplementary Results and Fig. S2 for training effects in specific load-sensitive PFC regions). Likewise, follow-up analyses in older adults confirmed a Time×Load cross-over interaction for loads 7-8 (*F*_1,18_=7.11, *p*=0.016, η_p_^2^=0.28), and a shift from a quadratic to a linear trend with training (see Supplementary Results). In task-negative regions, younger adults showed less deactivation with training (Time: *F*_1,20_=8.49, *p*=0.009, η_p_^2^=0.16), particularly for lower loads (Time×Load: *F*_4,80_=3.17, ε=0.59, *p*=0.043, η_p_^2^=0.14), whereas older adults showed no training effects (*p*s>0.6), consistent with evidence of less DMN modulation with aging (Turner & Spreng, 2015) (see Fig. 4b).

**Fig. 4.**
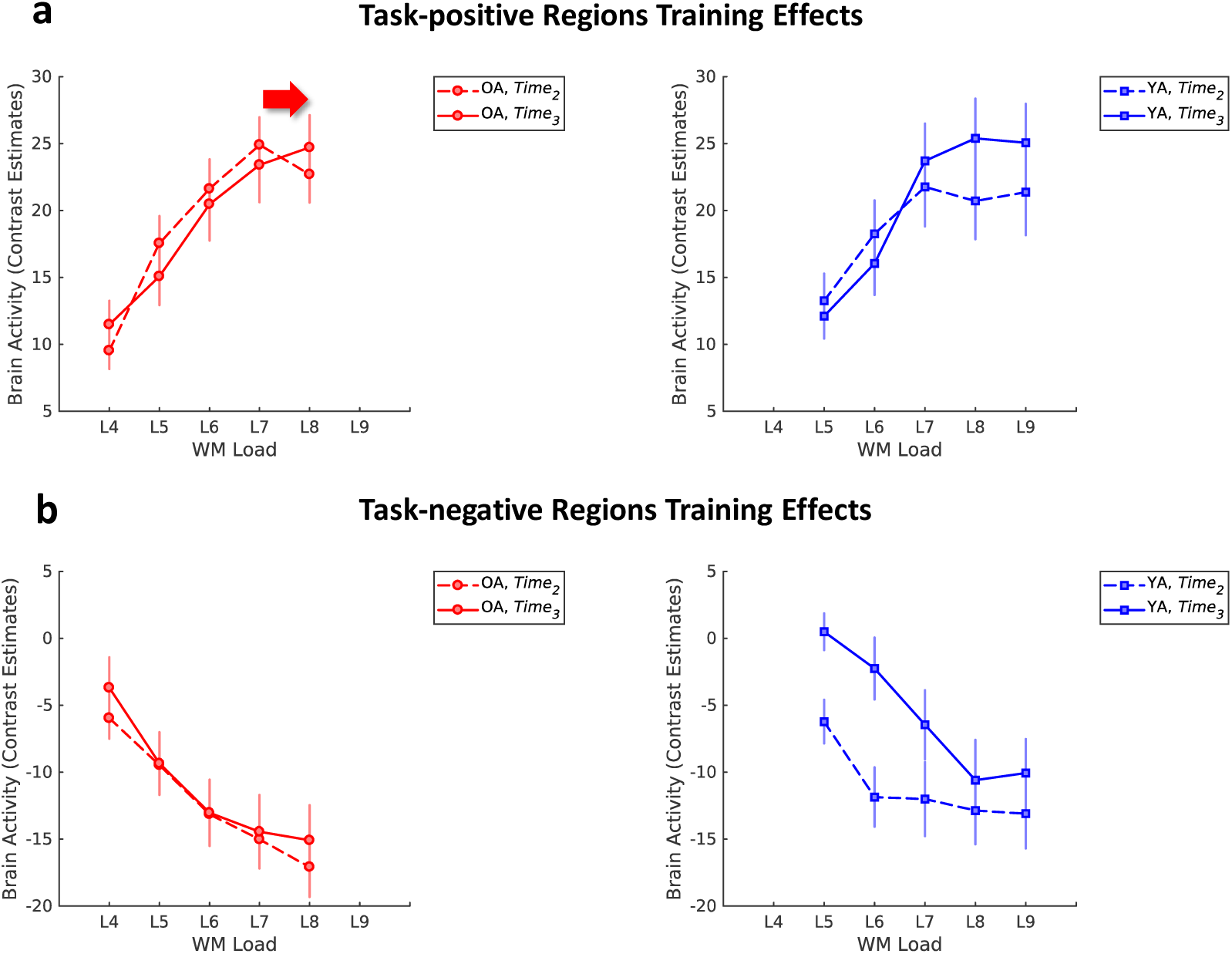
Training effects on brain activation in group-specific regions sensitive to load at Time1. **a**, Training effects in task-positive regions, associated with working memory. Both groups show greater activation at higher loads after training compared to before training. Older adults show a shift from quadratic to linear trends, consistent with the results in meta-analytically defined WM regions (see Fig. 3). **b**, Training effects in task-negative regions, associated with the default-mode network. Younger adults show less deactivation with training, particularly for lower loads. There are no training effects for older adults in these regions. Line-graphs display average brain activity for each load vs. baseline (load of 1) in group-specific regions sensitive to load at Time1 (see Materials and Methods and Fig. S1b). Error bars display standard error of the mean. WM, working memory; OA, older adults; YA, younger adults.

### Functional Connectivity Results

#### Task-exposure and Training Effects within the Meta-analytically Defined WM Network

Similar to the brain activation analyses, first we assessed effects of task-exposure and training on functional connectivity within the WM network. For this purpose, we employed the Power et al. (2011) functional atlas and selected those ROIs that overlapped with the Neurosynth meta-analytical WM map (23 ROIs; see Materials and Methods and Fig. S3a). We performed two Group×Time×Load ANOVAs on estimates of average functional connectivity (loads 5-8) within this meta-analytical WM network. The Time1 vs. Time2 analysis (Fig. 5, left panel) indicated no effects of task exposure on functional connectivity (all *p*s>0.15). Comparing Time2 and Time3 to assess training effects (Fig. 5, right panel) yielded a Time×Group interaction (*F*_1,38_=5.09, *p*=0.03, η_p_^2^=0.12), indicating increased within-WM network functional connectivity with training in older compared to younger adults. The main effect of Load was also significant (*F*_3,114_=3.57, *p*=0.016, η_p_^2^=0.09).

**Fig 5.**
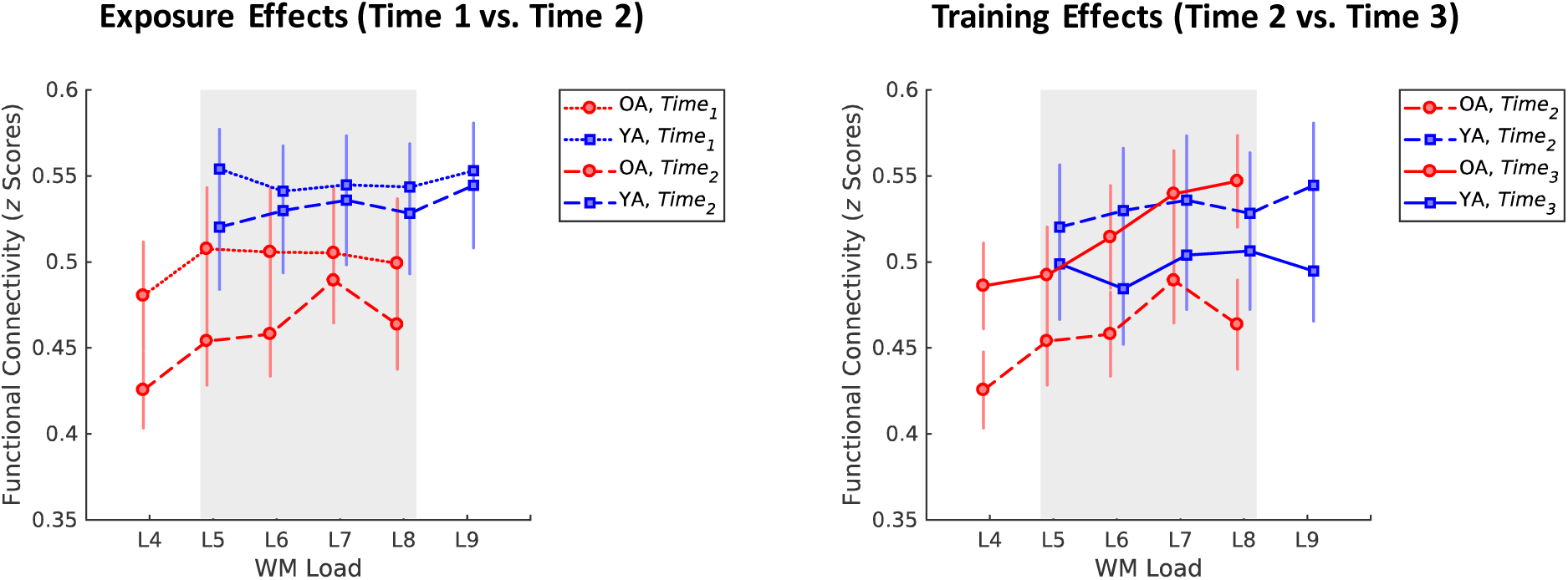
Exposure and training effects for functional connectivity within a meta-analytically defined WM network. Functional connectivity results show no significant differences pre-training (left panel) and increased functional connectivity within the WM network pre-vs. post-training, in older compared to younger adults (right panel). Line-graphs display average functional connectivity for each load vs. baseline (load of 1) within the meta-analytically defined WM network (see Materials and Methods and Fig. S3a). Error bars display standard error of the mean. The grey rectangle highlights the loads common to both groups (loads 5-8). WM, working memory; OA, older adults; YA, younger adults.

#### Training Effects within Group-specific Networks Sensitive to Load Manipulation

A second set of analyses examined training effects on functional connectivity, separately for each group, based on the Time1 data. First, task-positive and task-negative networks were defined based on the voxel-wise ANOVA results identifying brain-wide load-sensitive regions; network nodes were defined as peak-voxels within the task-positive and task-negative *F-*maps, separately for each group (see Fig. S3b and Tables 2 and 3 for employed coordinates). Then, training effects (Time2 vs. Time3) were analyzed separately for each group, using Time×Load ANOVAs across all loads (i.e., loads 4-8 in older adults and 5-9 in younger adults) on estimates of average functional connectivity within and between the task-positive and task-negative networks, respectively (see Materials and Methods).

Within the task-positive network, older adults showed greater functional connectivity with training (*F*_1,18_=9.41, *p*=0.007, η_p_^2^=0.34), thus replicating the results based on meta-analytical ROIs (see Fig. 6a; see also Supplementary Results and Fig. S5 for training effects on functional connectivity between specific load-sensitive PFC regions); younger adults only showed modulation of functional connectivity by WM load (*F*_4,80_=2.76, *p*=0.033, η_p_^2^=0.12). Within the task-negative network, older adults also showed a trend toward greater functional connectivity with training (*F*_1,18_=4.23, *p*=0.055, η_p_^2^=0.19), whereas younger adults showed no significant effects (*p*s>0.1) (see Fig. S4). Interestingly, older adults also showed greater functional connectivity between the task-positive and task-negative networks with training (*F*_1,18_=4.85, *p*=0.041, η_p_^2^=0.21), while both age groups showed modulation of between-networks connectivity with load (older adults: *F*_4,72_=4.58, *p*=0.002, η_p_^2^=0.20; young adults: *F*_4,80_=5.23, *p*=0.001, η_p_^2^=0.21) (see Fig. 6b; see also Supplementary Results for analysis of segregation between the task-positive and task-negative networks).

**Fig. 6.**
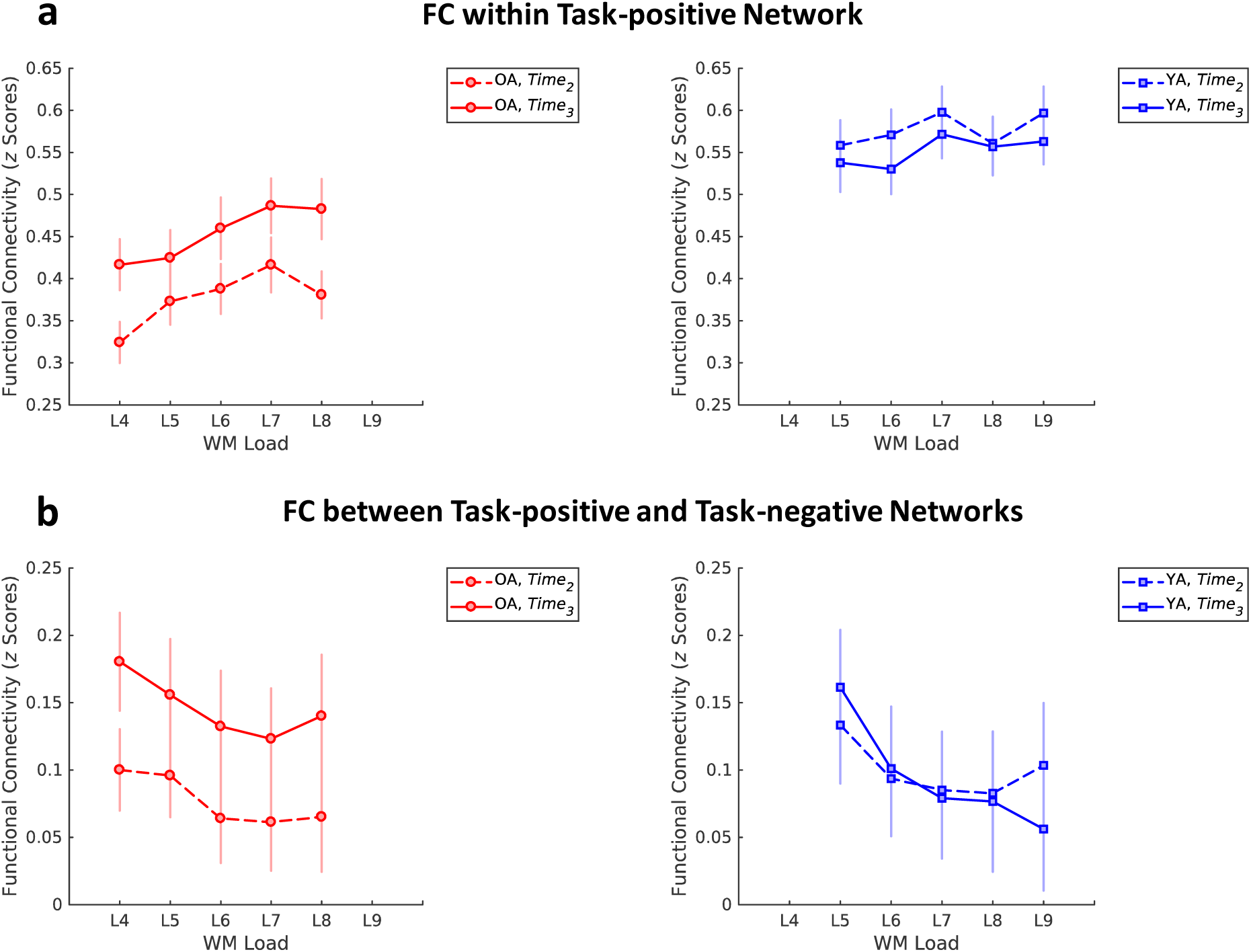
Training effects on functional connectivity in group-specific networks sensitive to load at Time1. **a**, Training effects on functional connectivity within the task-positive network, associated with WM. Older adults show greater functional connectivity with training, consistent with the results based on meta-analytical ROIs (see Fig. 5). Older adults also showed a trend toward greater functional connectivity with training within the task-negative network, which overlaps with regions of the default-mode network (see Fig. S4). There are no training effects in younger adults within either network. **b**, Training effects on functional connectivity between the task-positive and task-negative networks. Older adults show greater functional connectivity between the task-positive and task-negative networks with training. Line-graphs display average functional connectivity for each load vs. baseline (load of 1) within/between group-specific networks sensitive to load at Time1 (see Materials and Methods and Fig. S3b). Error bars display standard error of the mean. WM, working memory; OA, older adults; YA, younger adults.

## Discussion

CRUNCH posits that additional neural resources are recruited with increasing task demand regardless of age; however, older adults over-recruit at lower levels of demand compared to younger adults and reach capacity sooner with activation decline at higher loads. We hypothesized that WM training would bolster neural efficiency, lowering activation so that the activation peak shifts towards higher WM loads after training.

Younger adults performed better overall than older adults, but task-exposure reduced this age difference, and training improved performance for both groups. Brain imaging analyses yielded three main findings. First, older adults showed greater WM network recruitment pre-training, compared to younger adults, with activity decline at the highest load. Second, training led to greater brain responsiveness at higher loads for both groups, and for older adults, the activation peak shifted rightward toward higher loads. Finally, training increased task-related functional connectivity in older adults, both within the WM network and between this task-positive network and the task-negative/default-mode network. These results are discussed, in turn, below.

### Older Adults Over-recruit WM Regions and Show Activation Decline at High Loads

Using a meta-analytical WM map, we found greater recruitment of WM regions in older adults, with decline at the highest load, before training, and minimal effects of task exposure. Overall, these results are consistent with CRUNCH and replicate previous findings (Berlingeri et al., 2013; Cappell et al., 2010; Heinzel et al., 2014; Holler-Wallscheid et al., 2017; Kennedy et al., 2015; Schneider-Garces et al., 2010). According to CRUNCH, the extent of compensation-related activity varies with both the level of task demand and the resources available to meet that demand. For older adults relative to young adults, CRUNCH predicts over-activation for lower loads and under-activation or decline for the highest loads, as a function of both task demands and resource supply. Indeed, the present results support this prediction. Also in line with CRUNCH, the voxel-wise ANOVA at Time1 (see Fig. S1) showed bilateral modulation of brain activity by load in both groups, suggesting that contralateral recruitment is not unique to older adults, but an age-independent compensation mechanism engaged for difficult tasks (see also Holler-Wallscheid et al., 2017).

While here we replicated the quadratic activation trends previously identified in older adults, we did not find a group by load interaction, as was the case in our original investigation (cf. Cappell et al., 2010). Nevertheless, the activation slopes for older and younger adults at the highest loads had opposite signs (i.e., negative in older adults vs. positive in young adults), suggesting that a group by load interaction might have occurred had we included additional (higher) loads. These results could be partly explained by the inclusion of relatively young, high-performing older adults in the present sample (Moored et al., in prep.). Future investigations should consider inclusion of a more representative sample of older adults and use a larger range of loads capable of eliciting greater demand. Overall, though, the present pre-training results are consistent with CRUNCH which postulates age-related over-activation coupled with inefficient engagement at higher loads in older adults.

CRUNCH proposes that older adults compensate for neural inefficiency by over-activating at low loads to optimize task performance. Although the results comparing the two pre-training time points showed robust age differences in recruitment that cannot be simply explained by task exposure, a common factor, such as insufficient capacity or degraded neural function, could potentially explain both brain activity and decreased WM performance (see also Morcom & Henson, 2018) in older adults. In other words, based solely on pre-training data, one cannot assert a causal relationship between brain activation and behavior, because the evidence is simply correlational. In contrast, a within-subjects training intervention could potentially clarify the behavioral significance of neural over-activation (Iordan & Reuter-Lorenz, 2017; Lustig et al., 2009). Because CRUNCH makes clear predictions about the expected effects of training on brain activation (see Introduction) and given the within-subjects design of our study, a putative common factor could not explain both the pre- and post-training activation patterns, within the same individuals (see also Maillet & Rajah, 2013).

### Training Shifts Peak Activation in WM Regions toward Higher Loads in Older Adults

Our results show, for the first time, that adaptive WM training *increases* brain activity at higher memory loads, regardless of age, and shifts the demand-activation function in older adults. In general, more efficient networks are thought to exhibit lower activation than less efficient networks and can respond to a greater range of task demands (Barulli & Stern, 2013; Dunst et al., 2014). Assuming that brain networks become relatively less efficient with age, CRUNCH predicts both increasing activation with advancing age and narrowing of the dynamic range of activation (i.e., reduced ability to modulate brain activity in response to demand). While such predictions have been confirmed by previous cross-sectional investigations (e.g., Kennedy et al., 2015), the present results suggest that, to the extent that training improves efficiency, it also reduces the need for compensatory activation under low demand and enables greater activation under high demand (Festini, Zahodne, & Reuter-Lorenz, 2018; Lustig et al., 2009).

The present findings converge well with prior reports of age differences in task-related activation and training effects in older adults. First, these results are consistent with recent meta-analytical evidence (Li et al., 2015) showing that older adults’ propensity to over-activate the fronto-parietal control network is typically observed in studies in which older adults achieve performance levels comparable to younger adults (see also Maillet & Rajah, 2013). Second, our findings are also in line with previous training studies in older adults (e.g., Brehmer et al., 2011; Erickson et al., 2007; Heinzel et al., 2014) (reviewed in Duda & Sweet, 2019; Nguyen, Murphy, & Andrews, 2019). Such studies have typically identified decreased activation in WM regions after training, particularly for low and medium loads (e.g., 1- and 2-back conditions in an N-back WM task), but not for high loads (e.g., 3-back task) (Heinzel et al., 2014). Thus, these previous findings are consistent with a rightward shift of the neural recruitment curve with training, as predicted by CRUNCH, which can explain why both decreases and no changes (or even increases) (Duda & Sweet, 2019) in task-related BOLD activity have been reported in training studies. More specifically, either decreased *or* increased activation in task-relevant brain regions may occur when comparing pre-vs. post-training brain activity, depending on the level of demand elicited by the criterion task (i.e., the task employed to assess the training effects). This further underscores that parametric variation of task demands is critical for identifying boundary conditions (e.g., the crunch point) and that associations between brain activity and performance are likely non-linear (Reuter-Lorenz & Iordan, 2018).

The present evidence for training-related modulation of activity in fronto-parietal regions is in line with meta-analytical results suggesting that the most consistent loci of change with WM training are the same regions that are involved in WM performance (Constantinidis & Klingberg, 2016; Duda & Sweet, 2019; Salmi, Nyberg, & Laine, 2018). In addition to identifying training effects in fronto-parietal WM regions, however, we also identified less deactivation in task-negative regions with training in younger adults, particularly for lower loads. The task-negative regions overlap with the canonical default-mode network (DMN), which is anchored in the medial prefrontal and posterior cingulate cortices, and includes lateral parietal, as well as lateral and medial temporal regions (Buckner, Andrews-Hanna, & Schacter, 2008; Raichle et al., 2001). Typically, the magnitude of DMN deactivation increases with greater task difficulty (Mayer, Roebroeck, Maurer, & Linden, 2010; McKiernan, Kaufman, Kucera-Thompson, & Binder, 2003; Tomasi, Ernst, Caparelli, & Chang, 2006), presumably to allow maximization of resources for the WM/task-positive network. Thus, less deactivation in task-negative regions with training in younger adults suggests that the task has become easier, consistent with the idea of greater neural efficiency with training. The lack of DMN modulation with training in older adults is also in line with meta-analytical evidence from studies of brain activation (Duda & Sweet, 2019) and consistent with the idea of less modulation of DMN activity with aging (e.g., Turner & Spreng, 2012). In contrast, DMN modulation with training has been recently reported in younger adults (Finc et al., 2019).

### Training Increases Functional Connectivity within the WM Network and between the Task-positive and Task-negative Networks in Older Adults

Complementing the brain activation results, analyses of task-related functional connectivity showed both increased functional connectivity within the WM network and between the task-positive and task-negative networks in older adults, with training. Greater WM-network functional connectivity with training has been previously shown in younger adults with both resting-state (Jolles, van Buchem, Crone, & Rombouts, 2013; Takeuchi et al., 2013) and task-related data (Kundu, Sutterer, Emrich, & Postle, 2013), supporting the idea that strengthening of fronto-parietal coupling may benefit WM maintenance (reviewed in Constantinidis & Klingberg, 2016). Here, we observe a similar pattern, but only in older adults (see also Lebedev, Nilsson, & Lovden, 2018).

Although older adults showed increased functional connectivity within the task-positive/WM network with training, consistently for both meta-analytically defined and group-specific WM regions, these two analyses showed somewhat different profiles. Specifically, whereas the analysis using meta-analytically defined regions suggests opposite effects of training on functional connectivity within the WM network in older vs. younger adults (i.e., relative *in*crease in older adults vs. relative *de*crease in younger adults; see Fig. 5), the group-specific analysis conveys a general increase due to training in older adults and minimal effects in younger adults (see Fig. 6a). This indicates a tendency for older adults to be more responsive to training in our neural measures, a pattern we discuss further below.

The result showing increased functional connectivity between the task-positive and task-negative networks with training in older adults was not anticipated, given that the two networks are frequently described as being anti-correlated (Fox et al., 2005) and their competitive relationship is thought to be important for attention-demanding task performance (e.g., Kelly, Uddin, Biswal, Castellanos, & Milham, 2008). Recent studies, however, have challenged the idea that cognitive control reflects an antagonism between fronto-parietal and default-mode networks, and cooperation between the two systems has been detected in various cognitive tasks (see Cocchi, Zalesky, Fornito, & Mattingley, 2013). Thus, whereas the results based on the amplitude of brain activity may suggest that older adults become more “young-like” with training, the results based on task-related functional connectivity suggest complex and divergent age-related trajectories that remain to be further explored, considering both their potential benefits and costs (for instance, see Hillary & Grafman, 2017). The present results also underscore the complementary nature of activation-based and functional connectivity analyses.

The present functional connectivity results and to some extent, the brain activation results, suggest relatively more measurable functional reorganization with training in older adults than in younger adults. While such an outcome may initially seem surprising, it may be understood through Lovden’s theoretical framework regarding cognitive plasticity (Lovden, Backman, Lindenberger, Schaefer, & Schmiedek, 2010). Within this framework, flexibility refers to “the capacity to optimize the brain’s performance within the limits of the current state of functional supply” (p. 660), whereas plasticity relates to “the capacity for changes in flexibility” or “the capacity for changes in the possible range of cognitive performance” (p. 661). Because older adults presumably have a more limited range of functional supply, which in turn limits flexibility, the training intervention fostered greater plasticity (i.e., reorganization) in older adults. In other words, to the extent that the training intervention was overall more challenging for older adults, it may have promoted greater plasticity, which was reflected during the performance of the criterion fMRI tasks. In contrast, younger adults were potentially able to perform the tasks more within their range of flexibility, especially given the relatively short intervention. The currently employed paradigm, however, did not test the limits of training or plasticity in either group. Thus, future training studies with longer/more intensive interventions should further clarify whether behavioral and neural training effects persist over the long-term in the same task and whether they transfer to other working memory contexts. In addition, identification of boundary conditions, such as the crunch point of neural activation, could be further used to set personalized criterion tasks, whereas age-specific shifts of the demand-activation curves could potentially be used as neural markers of successful training interventions.

## Conclusions

In sum, the present results are consistent with CRUNCH and provide new evidence for the effects of training on brain activity as a function of age. According to the CRUNCH hypothesis, a resource ceiling that differs with age (Reuter-Lorenz & Cappell, 2008) but that can increase with training (Lustig et al., 2009), limits the system’s capacity to meet task demands. Consistent with this view, our results showed that older adults over-recruit WM regions and reach peak activation at lower loads than younger adults. As predicted, however, after training peak activation shifted to the right (to higher memory loads). This outcome suggests that appropriate training can increase the dynamic range of activation in WM circuitry (Kennedy et al., 2017), enabling greater responsiveness at higher loads. These results provide new evidence for functional plasticity with training in older adults and identify a potential signature of improvement at the neural level.

## Acknowledgements

This research was supported by a National Institute on Aging [R21-AG-045460] grant to P.A.R.- and a National Institutes of Health [1S10OD012240-01A1] grant. The authors thank Krisanne Litinas for assistance with MRI data reconstruction and Daniel Weissman for comments on a previous version of the manuscript.

## Author Contributions

P.A.R.-L., J.J., T.A.P., M.B., S.M.J., B.K., K.A.C., K.D.M., and S.J.P. designed the study. K.A.C. and K.D.M. collected the behavioral and brain imaging data. A.D.I. and K.A.C. analyzed the brain imaging and behavioral data. A.D.I. wrote the original draft. All authors reviewed and edited the final manuscript.

## Data Availability

The MRI and behavioral data that were used in this study are available to researchers from the corresponding authors upon request.

