## Supplementary Materials for "Neural Correlates of Working Memory Training: Evidence for Plasticity in Older Adults"

### Supplementary Results

***Robustness Analysis.*** Examination of distribution plots identified one older adult whose brain activation as a function of load was consistently greater than 1.5 times the interquartile range for all loads, at both Time1 and Time2. We repeated the analyses without this participant and the results did not change substantially. Specifically, regarding the exposure effects, the Time1 vs. Time2 analysis indicated greater overall recruitment in older adults ($F_{1,37}$=4.68, $p$=0.037, $\eta_p^2$=0.11). Regarding the training effects, the Time2 vs. Time3 analysis identified greater recruitment at higher loads with training, for both groups (Time×Load: $F_{3,111}$=3.63, $p$=0.015, $\eta_p^2$=0.09), and follow-up analyses confirmed a Time×Load cross-over interaction for loads 7-8 in older adults ($F_{1,17}$=14.83, $p$=0.001, $\eta_p^2$=0.47). Overall, this indicates that the observed effects were not substantially driven by this participant's data. There were no participants whose functional connectivity was consistently greater than 1.5 times the interquartile range.

***Multilevel Modeling of Brain Activity within Group-specific WM Maps.*** Similar to the results within the Neurosynth map, multilevel modeling to assess the influence of load on brain activity within the group-specific task-positive maps (see Materials and Methods) were consistent with a rightward shift of the CRUNCH curve only in older adults. Specifically, for older adults, a quadratic trend described the data over and above a linear trend before training (Time2: $-2LL_{\text{linear}}$=620.11, $-2LL_{\text{linear+quadratic}}$=589.74, $\chi^2_{\text{difference(1)}}$=30.38, $p$<0.001) but not after training (Time3: $-2LL_{\text{linear}}$=617.87, $-2LL_{\text{linear+quadratic}}$=614.63, $\chi^2_{\text{difference(1)}}$=3.24, $p$=0.072). In contrast, for younger adults, quadratic terms improved the model fit both before (Time2: $-2LL_{\text{linear}}$=732.29, $-2LL_{\text{linear+quadratic}}$=724.47, $\chi^2_{\text{difference(1)}}$=7.82, $p$=0.005) and after (Time3: $-2LL_{\text{linear}}$=729.92, $-2LL_{\text{linear+quadratic}}$=720.51, $\chi^2_{\text{difference(1)}}$=9.42, $p$=0.002) training.

***Training Effects in Load-sensitive PFC Regions.*** Because the Time1 maps were group-specific but not region-specific, we followed-up our main analyses by further examining training-related changes in two load-sensitive PFC regions identified at Time1, within these group-specific maps: the right dorsolateral PFC (dlPFC), a region involved in the executive control of WM and previously identified as a locus of compensatory support in older adults (Cappell et al., 2010), and the left inferior frontal gyrus (IFG), part of the verbal WM rehearsal circuit (Nee et al., 2013). We fitted 5 mm-radius spheres at the coordinates corresponding to the peak-activation voxels in each of these two regions at Time1, separately for each group (right dorsolateral PFC: MNI coordinates [x, y, z] = 36, 54, 21 in older adults and 36, 39, 33 in younger adults; left IFG: MNI coordinates [x, y, z] = -51, 6, 21 in older adults and -48, 9, 24 in younger adults; see Tables 2 and 3). The resulting contrast values for each participant, time point, and load were then exported to SPSS and similarly analyzed with repeated-measures ANOVAs (Time×Load), separately within each group. For the right dlPFC, older adults showed a trend toward reduced recruitment with training ($F_{1,18}$=4.14, $p$=0.057, $\eta_p^2$=0.19), whereas younger adults showed no training-related changes ($p$s>0.5) (see Fig. S2a). For the left IFG, older adults showed less recruitment for medium loads (Time×Load: $F_{4,72}$=3.16, $p$=0.019, $\eta_p^2$=0.15), whereas younger adults showed greater recruitment for higher loads (Time×Load: $F_{4,80}$=4.19, $p$=0.004, $\eta_p^2$=0.17), with training (see Fig. S2b).

Paralleling the supplementary brain activation analyses described above, we also examined training-related changes in functional coupling between the right dorsolateral PFC and the left IFG, using a similar repeated-measures ANOVA. Older adults showed no training effects and only a main effect of Load ($F_{4,72}$=5.06, $p$=0.001, $\eta_p^2$=0.22), whereas younger adults showed greater coupling with training, particularly for the load of 8 (Time×Load: $F_{4,80}$=2.75, *p*=0.034, $\eta_p^2$=0.12), and no main effect of Load (*p*=0.1) (see Fig. S5).

***Network Segregation.*** To summarize the interactions between the task-positive and task-negative networks with a single measure, we calculated their segregation (Chan, Park, Savalia, Petersen, & Wig, 2014), defined as the difference between within- and between-networks connectivity expressed as a proportion of within-network connectivity [i.e., *Segregation* = $(\bar{Z}_w - \bar{Z}_b)/\bar{Z}_w$, where $\bar{Z}_w$ is the within-networks connectivity and $\bar{Z}_b$ is the between-networks connectivity]. Results were analyzed with repeated-measures ANOVAs (Time×Load), separately within each group. There were no effects of training on segregation between the task-positive and task-negative networks, in either group, (*p*s>0.1) and only main effects of Load, in both groups (*p*s<0.001) (see Fig. S6). This suggests that despite the training effects on functional connectivity in older adults, both within and between the task-positive and task-negative networks, these changes were commensurate such that the ratio between within- and between-networks connectivity did not change.

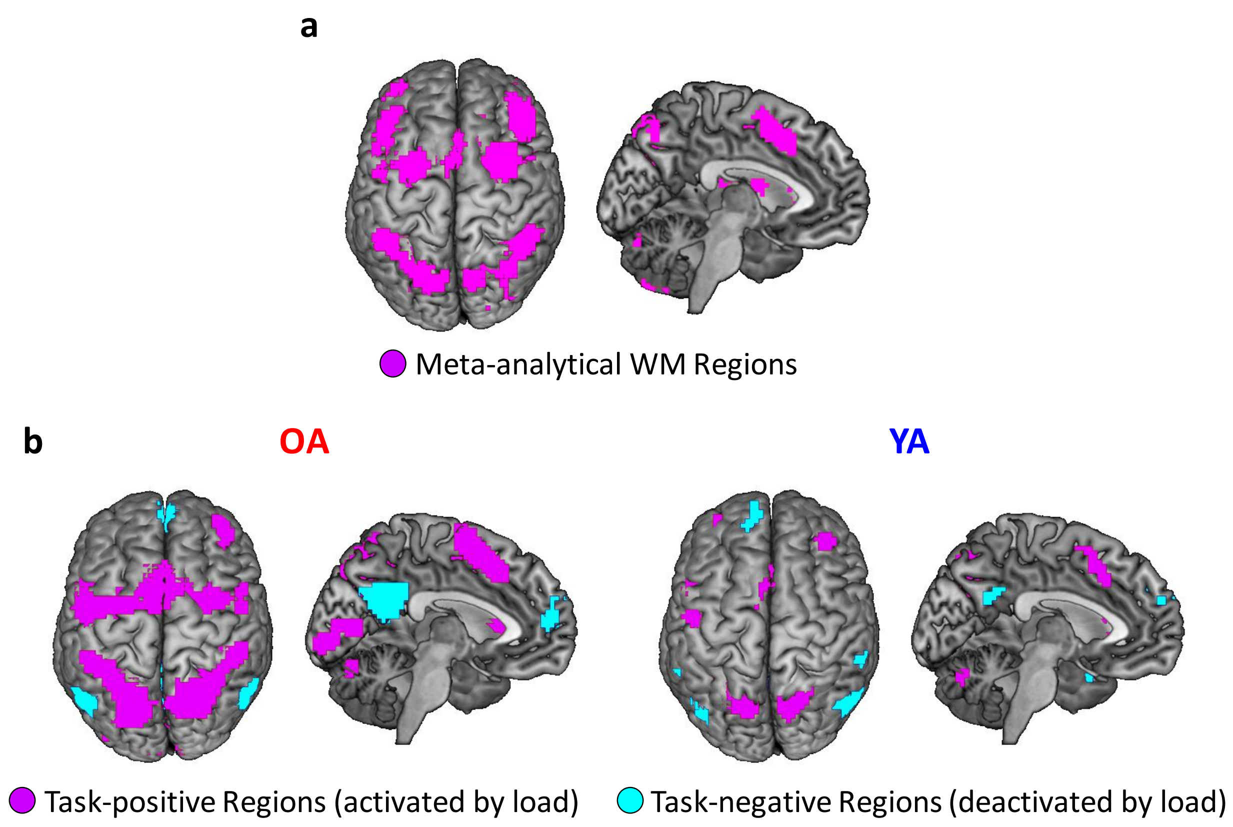


**Fig. S1. Brain regions involved in working memory and modulated by load. a**, Meta-analytically defined working memory (WM) regions, derived using Neurosynth (magenta map is thresholded at $p_{FDR} < 0.01$). **b**, Group-specific regions sensitive to load at Time1 (maps thresholded at $p < 0.001$, to convey the involvement of canonical default-mode network [DMN] regions in younger adults; brain activation analyses were performed using $p_{FWE} < 0.05$ corrected masks; see Materials and Methods and Tables 2&3). The magenta and cyan maps identify task-positive regions (i.e., *up*-regulated by load), associated with WM, and task-negative regions (i.e., *down*-regulated by load), associated with DMN, respectively. OA, older adults; YA, younger adults. Images were generated using MRIcron (http://mricro.com).

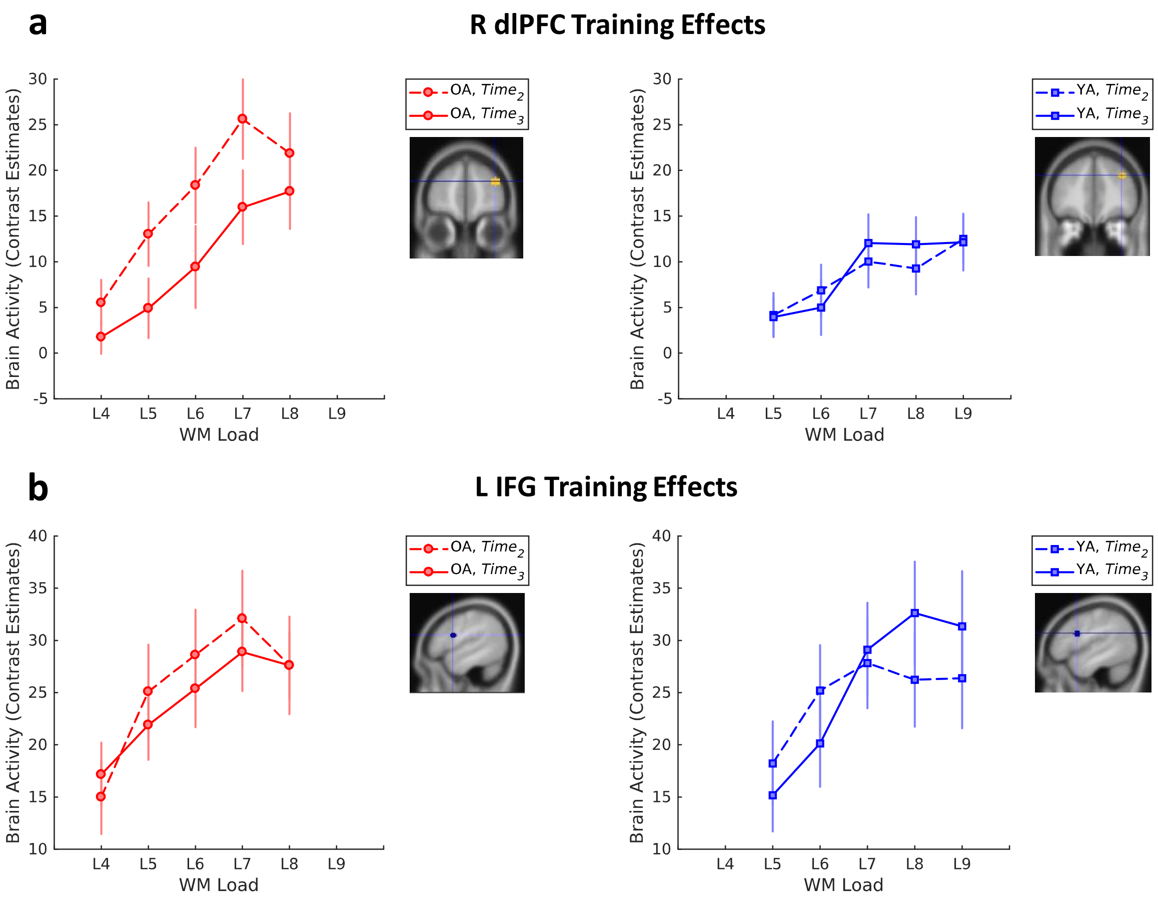


**Fig. S2. Training effects in task-relevant PFC regions, identified at Time1. a**, Training effects in the right dorsolateral prefrontal cortex (MNI coordinates [x, y, z] = 36, 54, 21 in older adults and 36, 39, 33 in younger adults). Older adults showed a trend toward reduced recruitment with training; there were no training-related changes in younger adults. **b**, Training effects in the left inferior frontal gyrus (MNI coordinates [x, y, z] = -51, 6, 21 in older adults and -48, 9, 24 in younger adults). Older adults showed less recruitment for medium loads, whereas younger adults showed greater recruitment for higher loads. Brain sections depict 5 mm-radius regions of interest fitted at peak-activation voxels at Time1 (see Tables 2&3). Line-graphs display average brain activity for each load vs. baseline (load of 1) within each region of interest (see Materials and Methods and Fig. S1b). Error bars display standard error of the mean. OA, older adults; YA, younger adults.

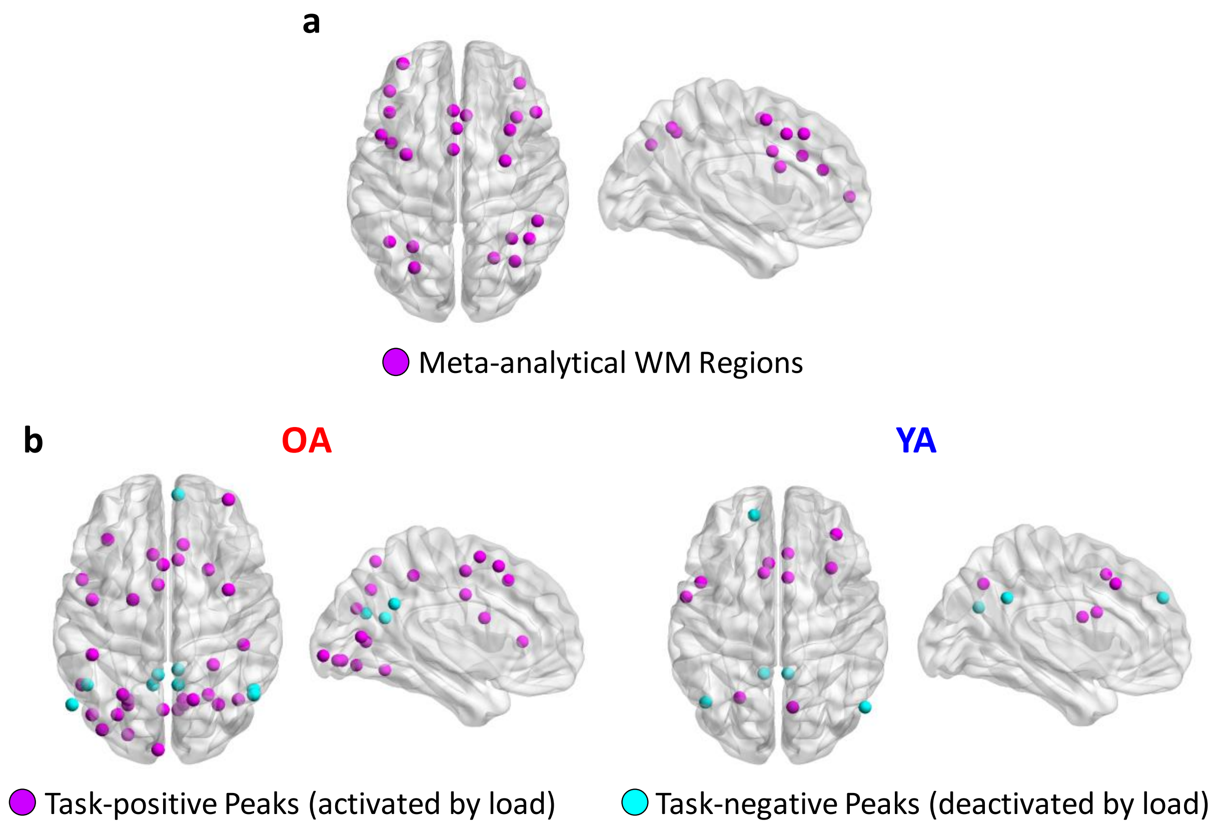


**Fig. S3. Brain networks involved in working memory and modulated by load. a**, Meta-analytically defined working memory (WM) network, derived using Neurosynth and the Power et al. (2011) atlas (see Materials and Methods). **b**, Group-specific networks sensitive to load at Time1. Each 5-mm sphere identifies the corresponding peak coordinate from Tables 2 and 3, respectively. The magenta and cyan maps identify task-positive regions (i.e., *up*-regulated by load), associated with WM, and task-negative regions (i.e., *down*-regulated by load), associated with DMN, respectively. OA, older adults; YA, younger adults. Images were generated using BrainNet Viewer (Xia et al., 2013).

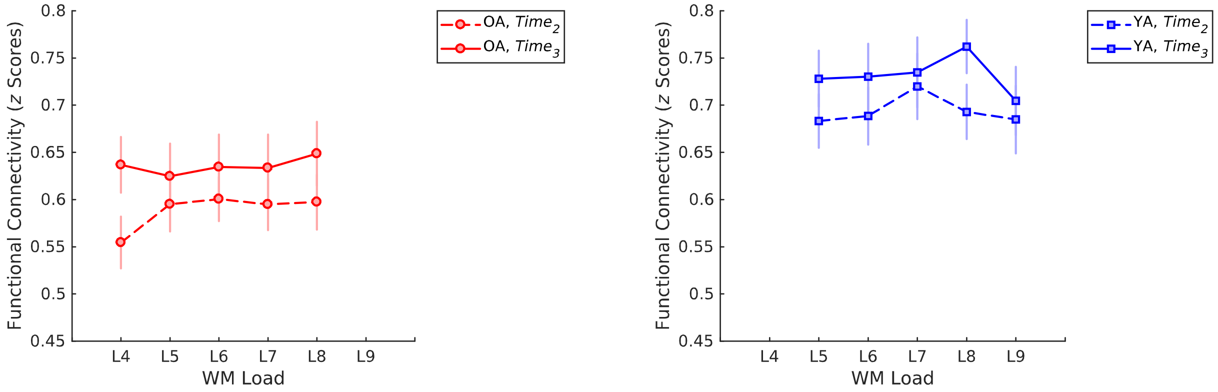


**Fig. S4. Training effects on functional connectivity within the task-negative network.** Older adults showed a trend toward greater functional connectivity with training, whereas younger adults showed no significant effects. Line-graphs display average functional connectivity for each load vs. baseline (load of 1) within the group-specific task-negative networks (see Materials and Methods and Fig. S3b). Error bars display standard error of the mean. OA, older adults; YA, younger adults.

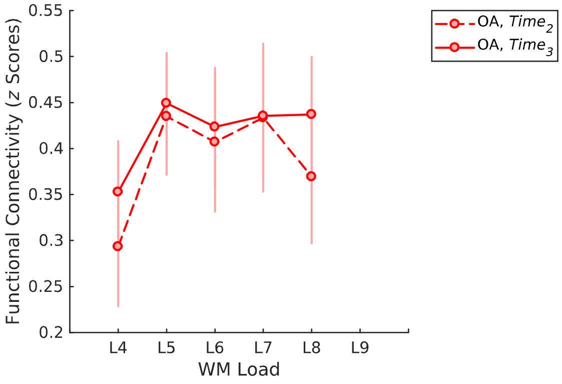


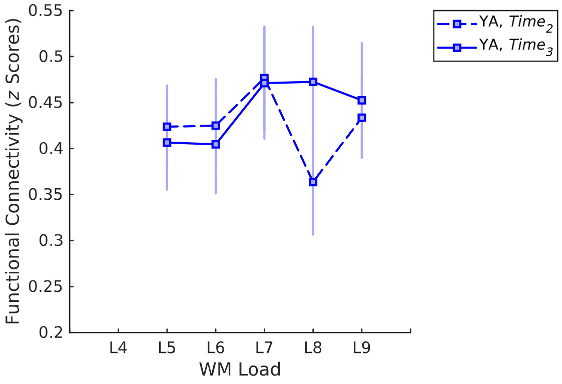


**Fig. S5. Training effects on functional connectivity between task-relevant PFC regions, identified at Time1.** Older adults showed no training effects, whereas younger adults showed greater functional connectivity with training for the load of 8. Line-graphs display average functional connectivity for each load vs. baseline (load of 1), between the right dorsolateral prefrontal cortex (MNI coordinates [x, y, z] = 36, 54, 21 in older adults and 36, 39, 33 in younger adults) and the left inferior frontal gyrus (MNI coordinates [x, y, z] = -51, 6, 21 in older adults and -48, 9, 24 in younger adults; see Tables 2&3). Error bars display standard error of the mean. OA, older adults; YA, younger adults.

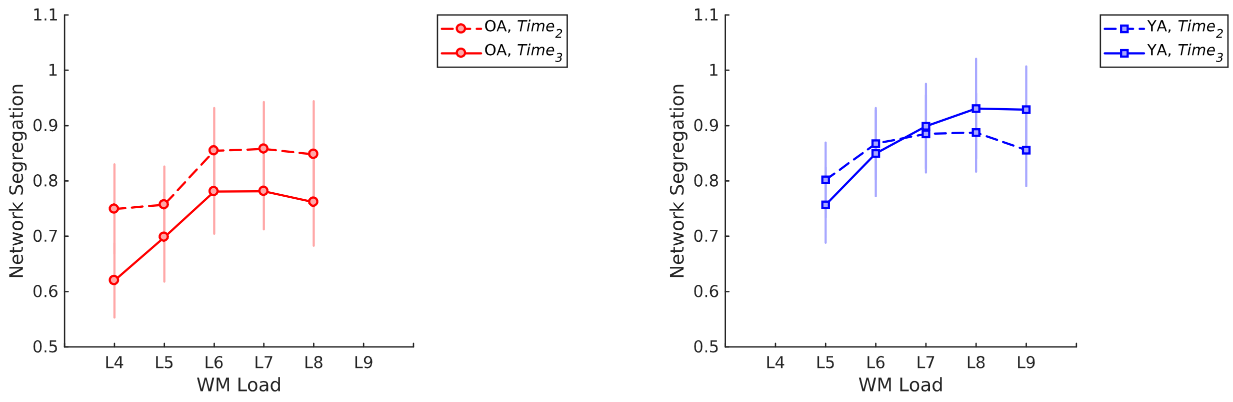


**Fig. S6. No training effects on segregation between the task-positive and task-negative networks.** Line-graphs display segregation for each load vs. baseline (load of 1) between the group-specific task-positive and task-negative networks (see Materials and Methods and Fig. S3b). Error bars display standard error of the mean. OA, older adults; YA, younger adults.
